# Dynamic Cognitive States Predict Individual Variability in Behavior and Modulate with EEG Functional Connectivity during Working Memory

**DOI:** 10.1101/2021.08.02.454757

**Authors:** Christine Beauchene, Thomas Hinault, Sridevi V. Sarma, Susan Courtney

## Abstract

Fluctuations in strategy, attention, or motivation can cause large variability in performance across task trials. Typically, this variability is treated as noise, and assumed to cancel out, leaving supposedly stable relationships among behavior, neural activity, and experimental task conditions. Those relationships, however, could change with a participant’s internal cognitive states, and variability in performance may carry important information regarding those states, which cannot be directly measured. Therefore, we used a mathematical, state-space modeling framework to estimate internal states from measured behavioral data, quantifying each participant’s sensitivity to factors such as past errors or distractions, to predict their reaction time fluctuations. We show how modeling these states greatly improves trial-by-trial prediction of behavior. Further, we identify EEG functional connectivity features that modulate with each state. These results illustrate the potential of this approach and how it could enable quantification of intra- and inter-individual differences and provide insight into their neural bases.

**Statement of Relevance:** Cognitive behavioral performance and its neural bases vary both across individuals and within individuals over time. Understanding this variability may be key to the success of clinical or educational interventions. Internal cognitive states reflecting differences in strategy, attention, and motivation may drive much of these inter- and intra-individual differences, but often cannot be reliably controlled or measured in cognitive neuroscience research. The mathematical modeling framework developed here uses measured data to estimate a participant’s dynamic, internal cognitive states, with each state derived from specific factors hypothesized to affect attention, motivation or strategy. The results highlight potential sources of behavioral variability and reveal EEG features that modulate with each state. Our method quantifies and characterizes individual behavioral differences and highlights their underlying neural mechanisms, which could be used for future targeted training or neuromodulation therapies to improve cognitive performance.

## Introduction

Identifying the neural underpinnings of working memory (WM), the ability to actively maintain and use task-relevant information (1–3), or any other cognitive process, requires identifying a relationship between the experimentally structured task rules and stimuli, and the generated neural and behavioral responses. Generally, these relationships are thought to remain consistent when identical task conditions (i.e. trial types and stimuli) are repeatedly administered over a session.

This is often not the case, however, because even within a set of identical task trials, large differences can exist in recorded brain activity (e.g. magnitude, neuroanatomical location) and behavior (responding faster or slower, correctly or incorrectly). Such variability can be observed both between participants, and within a single participant during a session. The conventional assumption is that variability within an individual in both neural and behavioral responses is a consequence of noise. However, a growing trend in cognitive science is the belief that spontaneous neural activity contains valuable information (4–6). In this study, we build upon these ideas and argue that variability within individuals may be attributed to factors that are not random noise, but which are also not directly observable or easily predictable, such as fluctuations in attention or motivation. In addition, each of these factors may be more or less influential on one individual’s behavior versus another’s.

For example, imagine various scenarios where such cognitive/motivational states might affect your response on a cognitive task. If you are consistently answering correctly, then you might feel more confident or motivated, resulting in faster response times in present and future trials. On the other hand, a streak of incorrect answers may cause you to become more cautious and slower in order to learn from your mistakes. The opposite may be true as well, in that you may instead get frustrated with a losing streak and therefore speed up your response time. In addition, you may choose a different strategy from another individual performing the same task, or you may change strategies as you learn more about the task demands. Such cognitive/motivational states can vary during a session. We refer to these short-term fluctuations that affect behavior as internal states, which are difficult to quantify because they are not directly measurable.

Internal states are reflected in neural activity, which may be an independent representation of the state or multiplexed with neural activity related to motor behavior or cognitive processes. Disentangling the motivational, behavioral, and cognition-related neural activation patterns is necessary for understanding how each state varies within and across individuals. Previous work by our group had shown that internal states could accurately predict variations in risk-taking in a gambling task (7–9).

We used a dataset from a previous study (10) in which a salient, distracting stimulus was presented during the delay period of a spatial working memory task (11, 12). Overall, individual reaction times (RT) varied greatly in all participants within a single session. In the current study, we used a state-space modeling framework to estimate internal states from measured behavioral data, which characterized the highly variable RTs recorded during a working memory task performed while electroencephalography (EEG) data were collected. Using the state-space model, we tested our hypothesis that RT is a function of the task, stimuli, noise, and *identifiable internal states*. We identified three internal states that significantly improved model performance. These states were related to the participant’s recent past performance (including previous errors and RTs), and to their previous susceptibility to distraction. By including these internal states in the behavioral model, the model performance improved by a factor of 10 and could reliably predict RT variability. We then performed a regression analysis to identify a set of EEG features that could predict the internal states.

Finally, we showed that we can substitute the estimated internal states using EEG features for the internal states derived from measured behavior to reliably predict RT variability from neural activity. We identified a small number of EEG features whose activity level directly correlated with the significance of the internal state to the prediction of RT. Specifically, we found that information regarding an individual’s accuracy on preceding trials is related to right fronto-parietal connectivity and left parietal local activity in the gamma band. An individual’s RT on preceding trials, on the other hand, is associated with left fronto-parietal connectivity in the gamma band and temporo-parietal connectivity in the alpha band. Finally, left frontal activity in the gamma band is linked with the effect of distractors on RT in past trials.

While results are in line with previous work on topdown control of information processing (13) and attention control (14), our findings suggest that variability in working memory performance within and across individuals is due to more than task parameters, cognitive capacity, and noise. Previous studies have also found effects of past performance and distraction on future performance in group analyses. The results here, however, demonstrated that the degree to which each cognitive internal state, and its associated EEG features, contributed to the prediction of RT was different for each participant. In addition, each cognitive state (and thus its contribution to RT) also varied across time within individuals. Therefore, the results suggest this approach could be used to better quantify and characterize individual participants’ differences in cognitive performance, their dynamics, and their underlying neural mechanisms.

## Methods

### Participants

In total, twenty-six participants were recruited to complete the task. Each participant reported normal or corrected-to-normal vision and gave written informed consent approved by the Institutional Review Board of Johns Hopkins University. In the original study by our group (10), we excluded seven participants from analysis because of excessive EEG artifacts (< 70% of epochs after preprocessing), and/or incorrect trials (< 60% accuracy on the task), and/or technical difficulties. For this work, we excluded an additional three participants because a substantial number of trials did not include recorded EEG data. As a result, the recorded data from sixteen right-handed participants (fourteen females, mean age 20.9 ± 2.8 years) was used in this analysis. Please refer to (10), for EEG ERP and time-frequency analysis of this dataset.

### Working Memory Task

In this study, we analyzed the behavioral and EEG data of 16 participants who completed a value-driven attention capture (VDAC) working memory (WM) task to test their ability to ignore a previously learned distractor, which was no longer relevant (10). Specifically, the task involved remembering one of two different types of spatial information (spatial location or relation between items) across a delay period. During the delay, irrelevant stimuli briefly appeared on both the left and right side of fixation, with a more salient, previously rewarded distractor being either on the same (congruent) or opposite (incongruent) side as the subsequent working memory test stimulus (Fig 1, for more detail refer to the Supplemental Material). Participants responded with a button press to indicate whether the test stimulus matched the remembered sample stimulus according to the relevant spatial dimension. In total, the participants completed 540 trials, where 270 Location trials and 270 Relation trials were pseudo-randomly presented.

**Fig. 1.**
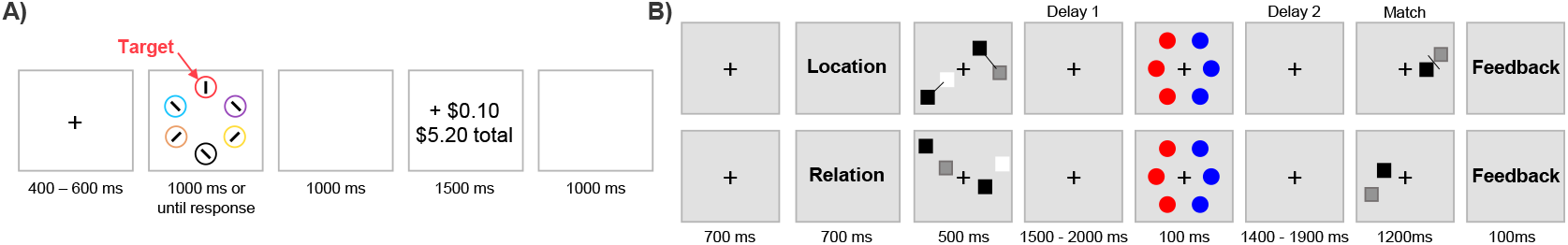
A) A schematic of the reward training task. B) Trial schematics showing the VDAC WM task for both the Location and Relation Trials, see (10) for additional details.

Additionally, 32 channels of EEG were recorded at 512Hz using an ActiCHamp System (Brain Products, Germany) and was referenced to Cz. The session took place in an electromagnetically shielded room to reduce noise.

### Developing Behavioral Models of RT

To predict RT using behavioral data for each participant, we used a general linear model (GLM). To assess the performance of the models, we computed the Pearson Correlation Coefficient, which is a measure of linear correlation between two variables (in this case, the predicted and recorded RT).

#### Pre-processing Behavioral Data

First, we interpolated the RT on trials (> 1%) where the participant timed-out (RT > 1200ms). Then, we filtered the RT timeseries by using a smoothing spline (smoothing factor of 0.15).

#### Model: Task + Noise (T + N)

We defined our initial model (Eqn 2) of RT to be a function of the Task Type, Hemifield, Congruent Trial, and Incongruent Trial. First, the Task Type, *s_t_*(*t*), was an indicator function where 0 and 1 represented the type of trial (location task vs. relation task), respectively. Next, the Hemifield indicator function, *s_h_*(*t*), identified the hemifield of the test stimulus (right vs left) on each trial. Finally, two separate indicator functions were used to specify the Congruent Trials, *s_c_*(*t*), and Incongruent Trials, *s_i_*(*t*).

We divided the trials into an interleaved training (80%) and testing set (20%). Using Matlab, we fitted the T + N model using a GLM to predict the RT, for each person. The resulting fitted GLM produced the *β*’s, and the associated *p*-value for each term of the model. Additionally, the *β*’s provided important information about how much that specific term contributed to the prediction of RT. Overall, we found this model didn’t capture the RT dynamics well.

#### Model: Task Type + Internal States + Noise (T + IS + N)

Based on the results of the initial model, we defined a modified model (Eqn 3) which included the initial T + N model, in addition to an exponential trend, and three internal states. Briefly, the internal states captured various aspects of the participants’ behavior and were generally defined as

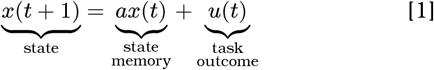

where the state, *x*, evolved over each trial (time step), *t*, by multiplying the current value of the state by a weight, *a*, which captured the contribution (or memory) of past task outcomes, *u*, to the current state value. Overall, state-space models were used to define the three internal states: the Previous Accuracy State (Eqn 4), Previous RT State (Eqn 5), and Previous Distraction State (Eqn 6).

##### Previous Accuracy State (x_1_)

the state received a positive unit input pulse on each trial that a participant answered incorrectly. Therefore, the state only deviated from zero if the participant made an incorrect response. If *a*_*x*_1__ was small (close to 0), then, on an incorrect answer, the state increased by one (due to the input pulse) and quickly decayed back to zero. However, if *a*_*x*_1__ was large (close to 1), then, again, the state increased in amplitude by one, and slowly decayed back to zero over the subsequent trials. If the state had not fully decayed to zero by the next incorrect answer, then the state could accumulate those recent errors. If the fitted weight (*β*_*x*_1__) from the behavioral model was positive, then, when the participant made an error, the state increased, and, as a result, the participant’s RT increased (or slowed down). The opposite occurred if *β*_*x*_1__ was negative, then, an error caused the state to decrease, which resulted in RT decreasing.

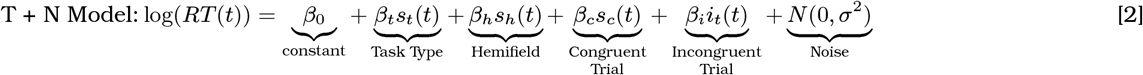

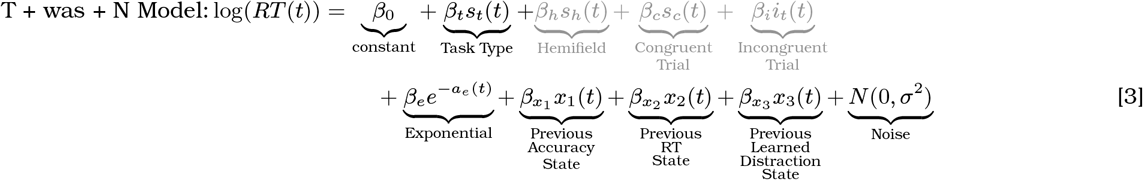

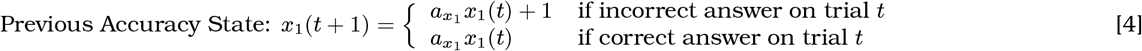

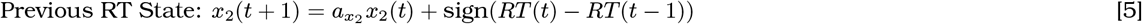

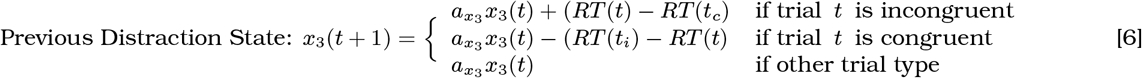

The T + N model tests four different experimentally structured task rules: Task Type, Hemifield, Congruent Trials, and Incongruent trials. Based on the results of the participant-specific fitted models, the Task Type term was significant (*p*-value < 0.05) in more than 50% of participants. Hemifield, Congruent trials, and Incongruent trials were significant in 25%, 0%, and 0% of participants, respectively. Therefore, to reduce the model complexity of T + IS + N, but still retain the task rules component, the Hemifield, Congruent, and Incongruent terms were removed (shown by the grayed out terms). For the Previous Distraction State, we defined *t_i_* to be the previous incongruent trial and *t_c_* to be the previous congruent trial. A congruent or incongruent trial was defined as when the two testing squares appeared in the same or opposite hemifield as the displayed rewarded distractor, respectively.

##### Previous RT State (x_2_)

a positive or negative unit input pulse was applied on each trial. The direction of the pulse was determined by the sign (positive or negative) of the difference between the RT of the current trial and the preceding trial. Therefore, the state accumulated information regarding the changing trend of the participants’ RT over time. If the fitted weight was positive, then the state fluctuated in the same direction as the RT. For example, if the participant was getting faster over the previous two trials (i.e. a negative input), then the state decreased as well, and RT decreased. Alternatively, if the fitted weight was negative, the opposite occurred and the trend was flipped (i.e. as the state decreased, RT increased).

##### Previous Distraction State (x_3_)

The purpose of this state was to quantify the difference in RT during trials where the previously rewarded distractor color was in the same hemifield (congruent) or opposite hemifield (incongruent) as the working memory test stimuli. This factor was previously associated with performance differences when averaging across trials and participants (10), but the magnitude of the effect of distraction varied both across participants and across trials within a participant. If a person continued to be affected by the distraction stimulus, then the “distraction” affected the congruent trials in the opposite direction as incongruent trials. For example, if a person’s attention was attracted by the distractor and the test stimulus was presented in the same hemifield (congruent trial), then RT decreased because their focus did not need to shift to the other side of the screen. However, if their attention was attracted by the distractor and then the test stimulus appeared on the opposite side (incongruent trial), then RT increased. This effect disappeared as the participants learned to ignore the distractor (i.e. delearned the previous association), and thus the predicted effect of congruence on the current trial converged to zero. For this state, an input pulse only occurred on congruent or incongruent trials. We defined the input to be the difference in the RT on the latest congruent vs. latest incongruent trial. The direction of this effect, however, was opposite depending on whether the current trial was congruent or incongruent. Specifically, the operator in front of the input depended on the current trial type, subtraction (shorter RT) for congruent or addition (longer RT) for incongruent trials. Similar to the Previous RT State, the sign of the fitted weight indicated the direction of the state fluctuations in relation to RT.

#### Behavioral model estimation and evaluation

We used a searchgrid to iterate through different combinations of *a* values (ranging between 0.01 and 0.9995) and fit the resulting GLM model (Eqns 3–6). To account for the gap in working memory task blocks when retraining the distractor color (after trials 180 and 360), we reset each of the three states back to zero on those trials. For each search grid iteration, we used the same training (80%) and testing (20%) set.

To find the final model, the minimum RMSE value (computed between the predicted RT and the RT training dataset) over all iterations was selected. Then, using the selected final combination of *a*’s, the GLM was fit to identify the *β*’s, but in this iteration, each of the terms of the model were normalized to be able to compare across participants. Overall, this process was repeated sixteen times to identify participant-specific behavioral models.

### Developing Neural Models of RT

Next, we developed a neural regression model to identify a set of EEG features that could predict each of the internal states and RT dynamics, for each participant. For additional details, please refer to the Supplemental Material.

#### Pre-processing EEG Data

For each of the 540 trials per person, the raw EEG data were pre-processed using the EEGLAB toolbox (version 14.4.2b) (15) in Matlab2020a. First, the data were bandpass filtered (1650^th^ order FIR filter) between 1Hz – 55Hz, and then re-referenced to the average. Next, the mean value of each channel was subtracted. Subsequently, automatic continuous rejection was applied using EEGLAB’s default independent component analysis algorithm (16, 17). Finally, the clean EEG was bandpass filtered (1650^th^ order FIR filter) into the five different frequency bands: Theta (4Hz – 8Hz), Alpha (8Hz – 12Hz), Low Beta (12Hz – 20Hz), High Beta (20Hz – 30Hz), and Low Gamma (30Hz – 55Hz).

#### EEG Feature Extraction

Using the cleaned EEG data, for each trial, we computed five pairwise Phase Locking Value (PLV) matrices, one per frequency band. PLV measured the phase coherence between two signals (18). For this analysis, we computed the PLV matrices from the EEG data 1400ms prior to the distractor presentation (during Delay 1), because this epoch could be considered to be the working memory maintenance portion of the trial.

Using the PLV matrices that were computed for each frequency band, on each trial, for each person, we extracted the relevant features of each matrix. To reduce the number of features, we defined seven regions of electrodes, which were left–frontal (Fp1, F7, F3), right–frontal (Fp2, F4, F8), central (FC1, FC2, CP1, CP2), left–temporal (FT9, FC5, T7, C3, TP9, CP5), right–temporal (FC6, FT10, C4, T8, CP6, TP10), left–parieto–occipital (P7, P3, O1), and right–parieto–occipital (P4, P8, O2).

In general, we were interested in Within Region and Between Region features. Specifically, to compute the Within Region features, we averaged all the PLV connections within each of the regions. For the Between Region features, the computation was similar except now we computed the average connection between regions. To compare the neural models across participants, we normalized and applied a smoothing spline (smoothing factor = 0.15) to the EEG features.

#### Neural Model and Feature Selection

The purpose of the neural model was to predict the internal states by using a combination of EEG features. We applied an additional GLM which was defined as

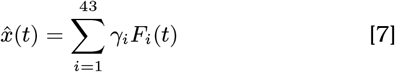

where 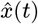 was the normalized estimated internal state on trial *t*, and *γ_i_* was the weight multiplied by each of the EEG feature vectors, *F_i_*(*t*).

To prevent overfitting, we used only 10% of the number of trials in the training dataset, which was 43 EEG features. Therefore, we applied a feature selection methodology to identify groups of EEG features for each state that were consistently significant across participants. Using the selected EEG feature sets, we computed a participant-specific neural model to predict each internal state.

### Predicting RT using estimated internal states from EEG features

To determine if RT could be predicted using EEG, we substituted the estimated internal states, using the EEG features, from the neural model (Eqn 7) for the original internal states (Eqn 4, 5, 6) of the behavioral model. Then, we re-fit the GLM for each person and computed the correlation values between the predicted RT and RT.

### Identifying EEG features weights which modulate with the magnitude of the internal state weights for predicting RT

As a final analysis, we focused only on the fitted weights (*β* and *γ*), which were fixed, constant, parameters in the participant-specific models. We identified the strongest linear relationships between the fitted weights of the internal states from the behavioral model (*β_x_*) and the weights of the EEG features in the neural model (*γ*).

The following process was completed for each internal state, but, for this example, we used the Previous Accuracy State, *x*_1_. For Participant 1, we plotted the absolute value of the Previous Accuracy State weight, |*β*_(*x*_1_,*P*_1_)_|, from their behavioral model against the absolute value of the neural model weight for the first selected EEG feature, |*γ*_(*F*_1_,*P*_1_)_|. This resulted in one point in the |*β*_*x*_1__| vs |*γ*_*F*_1__| space. Then, we repeated this process for all participants, *P*. Next, we fitted a linear model to the 16 data points, which was generally described by

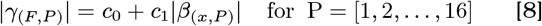

where *x* = [1, 2, 3] was the state, *F* = [1, 2,…, 43] was the EEG feature, and *P* was the 16 participants. The intercept and slope of the regression line were *c*_0_ and *c*_1_, respectively. From the linear model, we extracted the *R*^2^ value. We repeated this process for all 43 EEG features, for each state, and identified which neural features modulated with the contribution of the states to the RT prediction. We identified seven EEG features in which, across all 16 participants, the weights determined from the neural model strongly correlated (*R*^2^ > 0.4) with the weights of the internal states determined from the behavioral model.

## Results

### Reaction time is highly variable during a working memory task

In this study, the reaction time (RT) varied not only between participants but also within a single participant across trials (Figure 2A). Overall, the RT time series showed the characteristic exponential decrease in RT, potentially reflecting learning of the task. Also, the high-frequency oscillations observed were hypothesized to have been due to stimulus differences across trials, noise, or latent variables correlated with motivation, attention, or confidence. A summary of the variation in mean and standard deviation of the RT time series over the different participants is displayed in Fig 2B. The mean accuracy (percentage of correct answers over the 540 trials) and the mean RT, for each participant, are shown in Fig 2C.

**Fig. 2.**
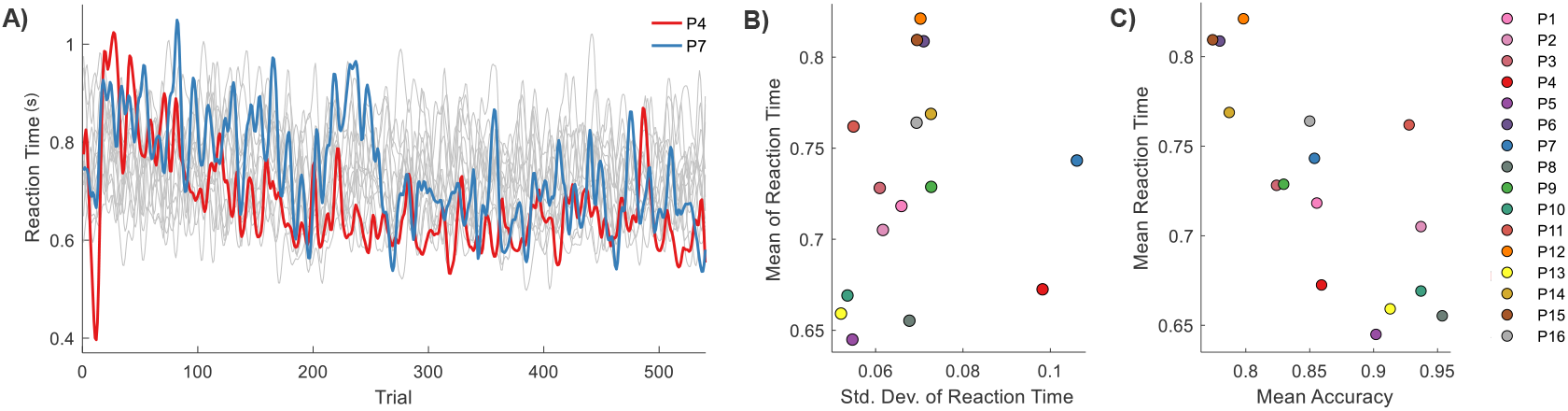
A) RT time-series traces for all participants. Participant 4 (red) and Participant 7 (blue) are highlighted. B) A comparison of the mean and standard deviation of the RT, for all participants. C) A comparison of the mean task accuracy, and mean RT over all participants.

### Dynamic internal states reliably predict trial-by-trial reaction time across participants

We first asked: “Can the observed highly variable reaction times be predicted with a model using only experimentally structured task rules and noise?” Based on the fitted parameters of the model (Eqn 2), we found the Task Type (location or relation trials) to be the most significant behavioral model term for predicting trial-by-trial RT. This was consistent with the previous work by our group on this dataset (10), which showed in a hypothesis-driven analysis that the RT was significantly shorter for location than relation trials, across trials and participants. However, the T + N model (Fig 3A) could not accurately capture the overall dynamics observed in the RT: Correlation (Mean ± SD): 0.07 ± 0.09.

**Fig. 3.**
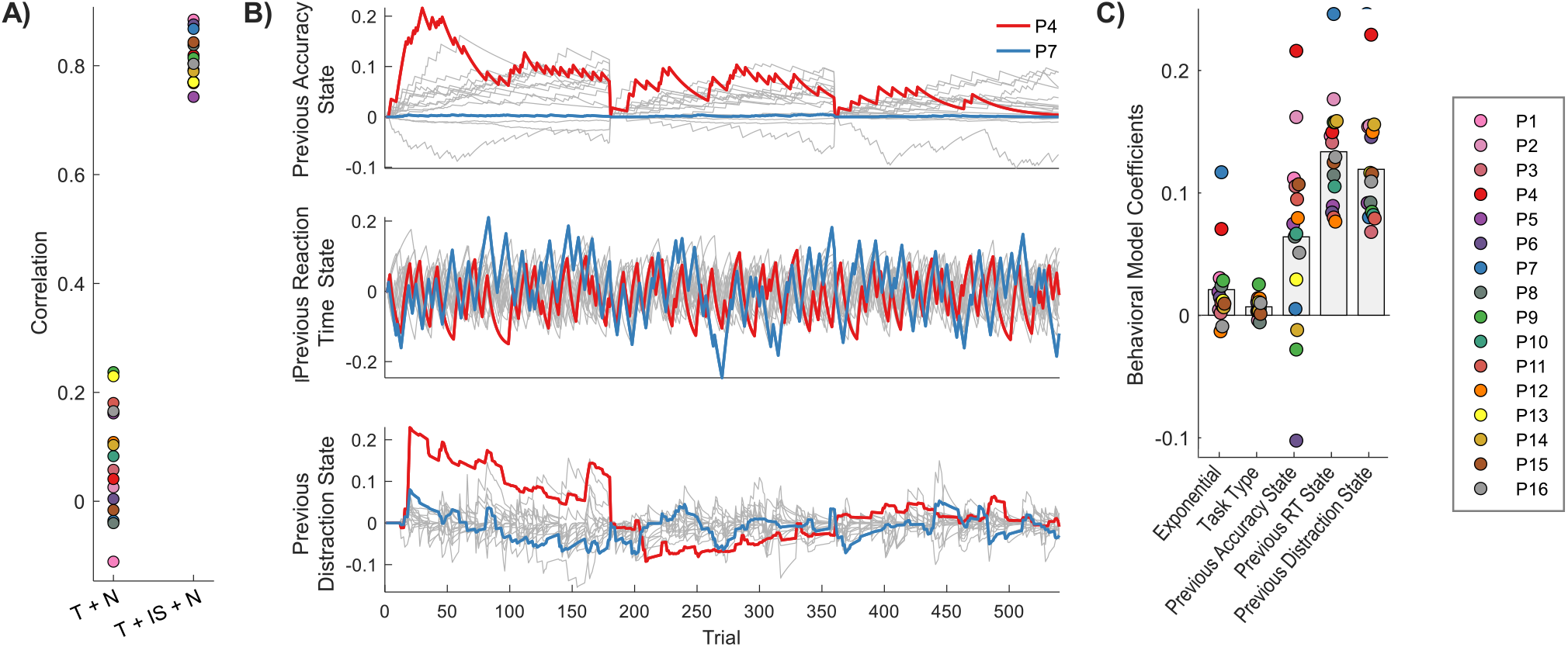
A) A comparison of the testing dataset correlations for the two models. T = Task, IS = Internal States estimated from behavioral model, N = noise. B) Trajectories for each of the states, for all participants. The states are reset to 0 at trial 180 and 360 to account for breaks that occurred between task blocks. Participant 4 and Participant 7 are highlighted in red and blue, respectively. C) The fitted *β*’s of each model term, for all participants.

Therefore, we next asked “Can internal cognitive states, in addition to task rules and noise, predict reaction time?” As shown in Fig 3A, we could accurately capture each participant’s RT time series with the T + IS + N model by using the three internal states, Task Type, exponential decay, and noise: Correlation (Mean ± SD): 0.81 ± 0.04).

Using the fitted participant-specific dynamical models, we uncovered which terms of the model (Eqn 3) weighed the most into predicting RT, for each participant. The trajectories for each of the states (scaled by their respective *β*’s) are shown for each person in Fig 3B. We chose to highlight Participant 4 and 7 throughout this analysis because they had similar accuracy and large RT variability, but they weighted their internal states very differently.

Overall, the participants varied in which terms were weighted (*β*) the most in their respective models (Fig 3C). Across all participants, the Exponential weight was generally positive, resulting in a negative exponential trend, suggesting task learning. In addition, the Task Type (Location or Relation working memory) resulted in the smallest weights across all participants but was still a significant model term in five participants. The weights of the internal states were generally larger than the weights associated with the other model terms (not including the constant term: −0.44 ± 0.25).

The Past Accuracy State weights were generally positive, which showed that as the participant accumulated errors, their RT increased (participants were slower to respond). Only in three participants were the fitted weights negative, which showed the opposite trend occurred, where they responded faster with accumulated incorrect answers. In addition, all weights for the Previous RT State were positive, which suggested that the positive or negative RT trend over the previous two trials continued on the current trial. Finally, for the Previous Distraction State, all the weights were positive, so the positive or negative difference between the previous incongruent and congruent trials was always applied in the same direction but was scaled by the weight.

Two examples of the participant-specific models are displayed in Fig 4A, illustrating how the three internal states differentially affect RT in individual participants. The top panel shows the fitted exponential curve and the indicator function for the Task Type. The middle panels show the three internal states’ evolution over the 540 trials. Finally, the bottom panels show the RT overlaid with the predicted RT from the model. To better see the model prediction of RT, Fig 4B shows scatter plots for the two participants, separately showing the data points used for training and testing the model. Overall, we fitted a participant-specific dynamic state-space model which could successfully capture a large portion of the variability of the RT for all participants, even though the factors driving RT could vary in each individual.

**Fig. 4.**
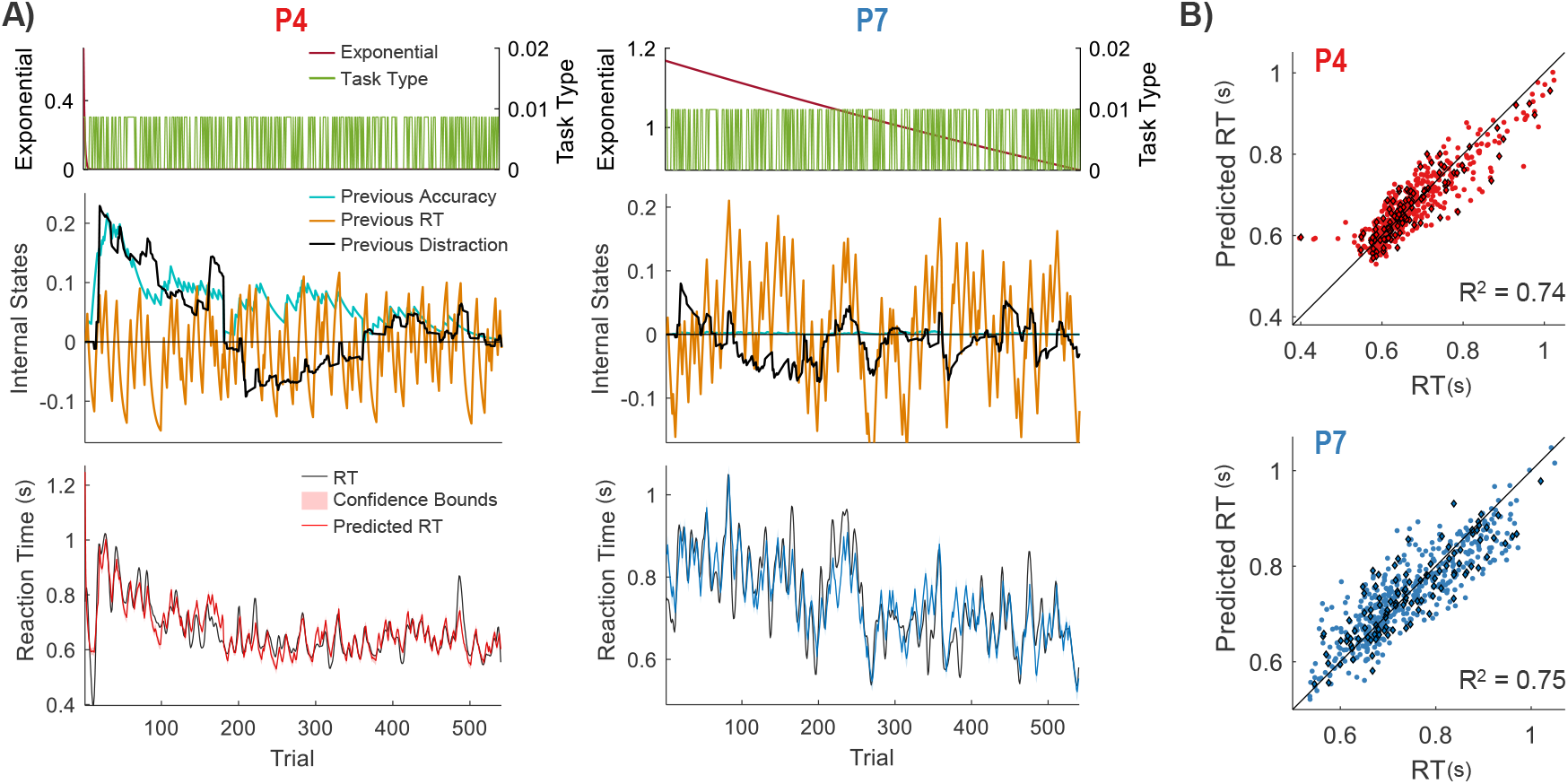
A) The top panels show each term of the model (scaled by its respective *β*). The exponential for P4 decays quickly. The bottom two panels show the predicted RT from the model overlaid with the RT. B) A scatter plot comparing the RT to the predicted RT. The training and testing data points are denoted by the colored circles and colored diamonds outlined in black, respectively. The solid black line shows perfect correlation (i.e. the *y* = *x* line)

### Groups of EEG features relate to each internal state

Using the estimated internal states identified for each participant, we asked “Does activity in groups of EEG features relate to each internal state?” Therefore, we generated a neural model (defined by a GLM) using the EEG features to predict the internal states. Fig 5A shows the percentage of the 43 selected EEG features selected from each of the five frequency bands, for each state. Overall, the largest number of selected features was found in the gamma band. Fig 5B shows, for each state, the EEG features that were significant (*p*-value < 0.05) in over 65% of the participant-specific neural models. Below the EEG features for each state is a scatter plot, for all participants, comparing the predicted internal state estimated from the neural model of EEG features to the corresponding state estimated from the behavioral model.

**Fig. 5.**
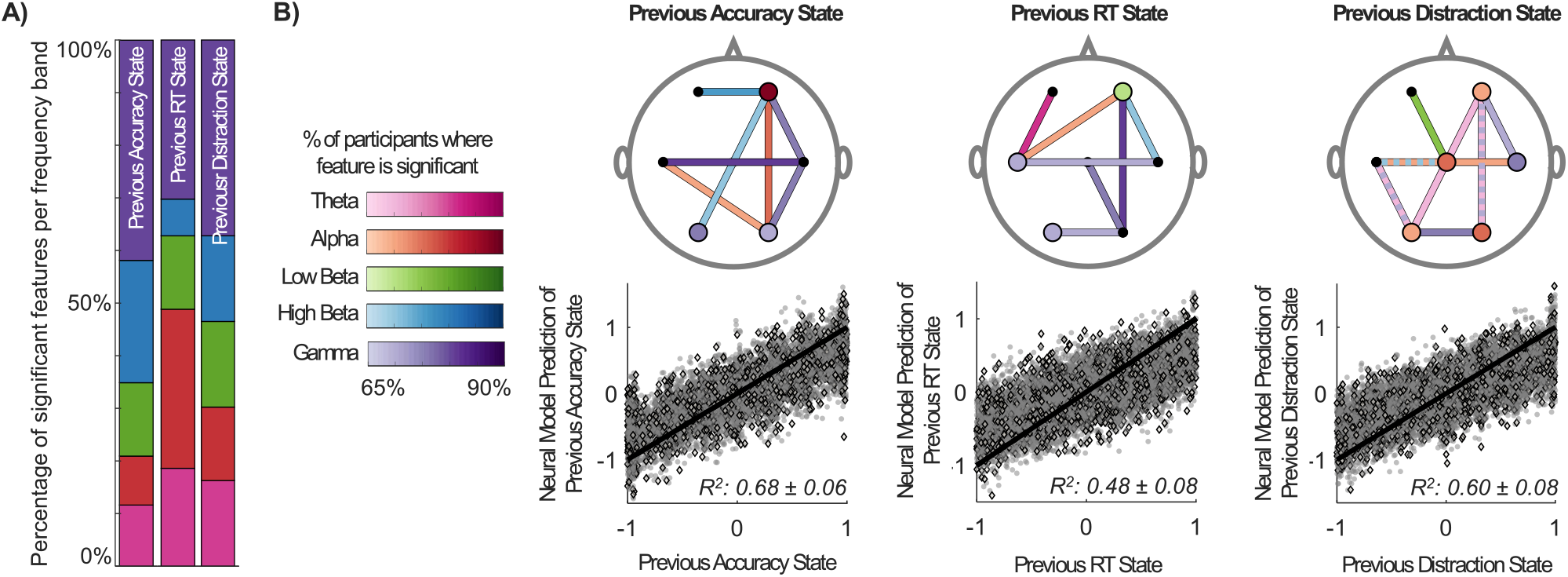
A) Distribution of the features selected in each frequency band for each internal state. The color to denote each frequency band is the same as in B (Theta is pink, Alpha is red, Low Beta is green, High Beta is blue, and Gamma is purple). B) The features that were significant in over 65% of participants. A feature was considered to be significant if the resulting feature-specific *p*-value was less than 0.05 from the GLM (i.e. *β* was significantly different from zero). The plots are shown in the transverse plane and the hemispheres are not flipped. The color of the feature shows the percentage and the frequency band. The Within Region features are shown as colored circles with a black outline. The Between Region features are shown as a colored line with two small black dots at either end. A dotted line shows the feature is present in two different frequency bands. The scatter plots show the performance of the neural model in predicting the internal states. The training and testing data points are denoted by the grey circles and grey diamonds outlined in black, respectively. The solid black line indicates perfect correlation (i.e. the *y* = *x* line)

### RT can be predicted using EEG features

Next, we asked, “Can RT be predicted using EEG features related to the internal states?” By substituting each of the internal states estimated from the EEG features into the behavioral model, we were able to capture the trial-by-trial RT dynamics, for each person. Fig 6A shows both the time series responses and scatter plots of the RT compared to the predicted RT using the EEG features, for Participants 4 and 7. Fig 6B shows how the model performance, quantified using correlation, changed if we substituted our internal states from the behavioral model with the estimated internal states using the EEG features (IS(EEG)). Over all participants, the drop in model performance was small and therefore showed that internal states could be related to neural activity measured using EEG. Furthermore, these results showed that this activity could predict a large portion of variability in RT, both within and across individuals.

**Fig. 6.**
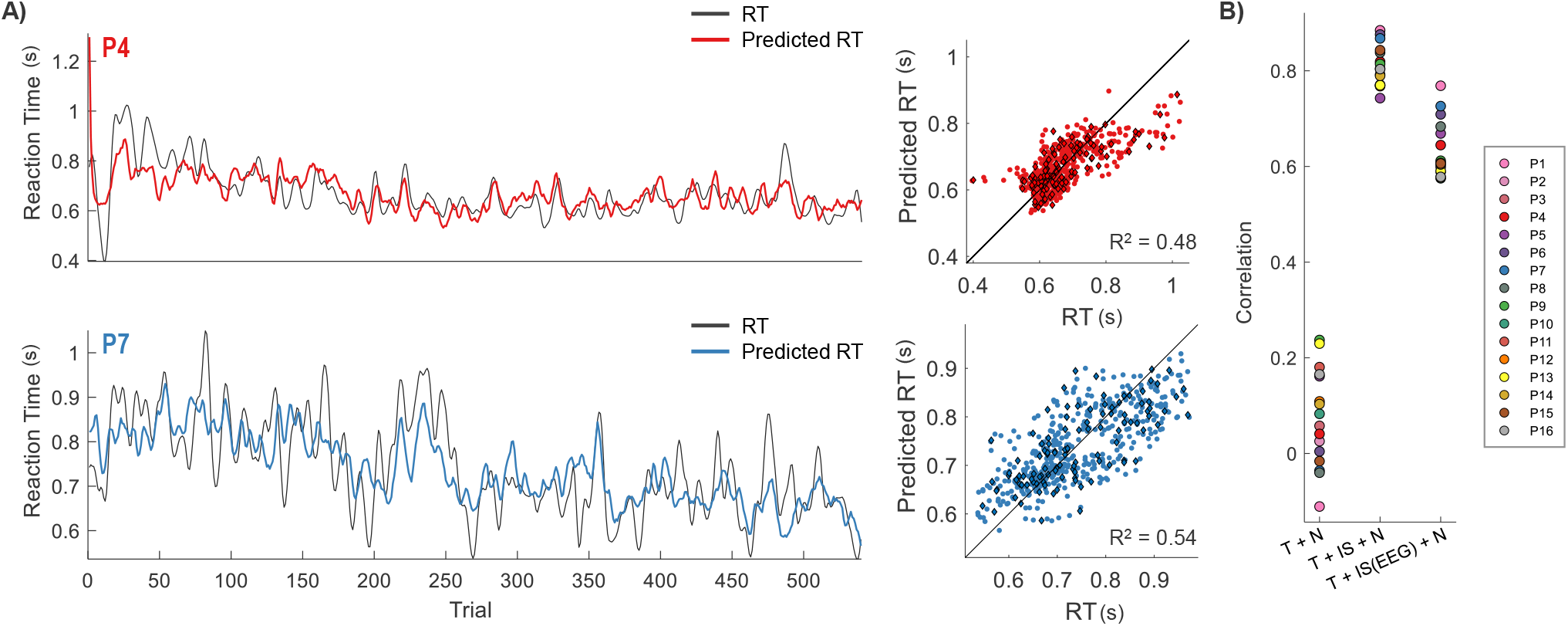
A) A comparison of the RT and the predicted RT using the estimated internal states from the EEG features. For the scatter plots, the training and testing data points are denoted by the colored circles and colored diamonds outlined in black, respectively. The solid black line shows perfect correlation (i.e. the *y* = *x* line) B) An extension to Fig 3A which shows the testing correlation for the dynamical model (T + IS + N) when the states were estimated using the EEG features (T + IS(EEG) + N). T = Task, IS = Internal States estimated from behavioral model, IS(EEG) = Internal States estimated from neural model, N = noise.

### Weights of individual EEG features modulate linearly with the weight of the internal state for predicting RT

Finally, we asked “Are there correlations between the participant-specific weights of a state in the behavioral model and the weights of a corresponding EEG feature in the neural model?” Specifically, we identified seven EEG features in which, across all 16 participants, the weights determined from the neural model (|*γ*|) strongly correlated (*R*^2^ > 0.4) with the weights of the internal states determined from the behavioral model (|*β*|).

The plots shown Fig 7 can be interpreted based on the direction of the correlation. If the correlation was positive, then, as the internal state weighed into the behavior more (i.e. increase in weight along the x-axis), the EEG feature weight for the internal state prediction also increased (i.e. increase in weight along the y-axis). However, if the correlation was negative, then, as the internal state weighed into the behavior more, the opposite occurred and the EEG feature weighed into the internal state less. Generally, across all three internal states, we observed a positive correlation between the contribution of the internal states in predicting RT, and the resulting magnitude of the EEG feature weight from the neural model for predicting the internal state.

**Fig. 7.**
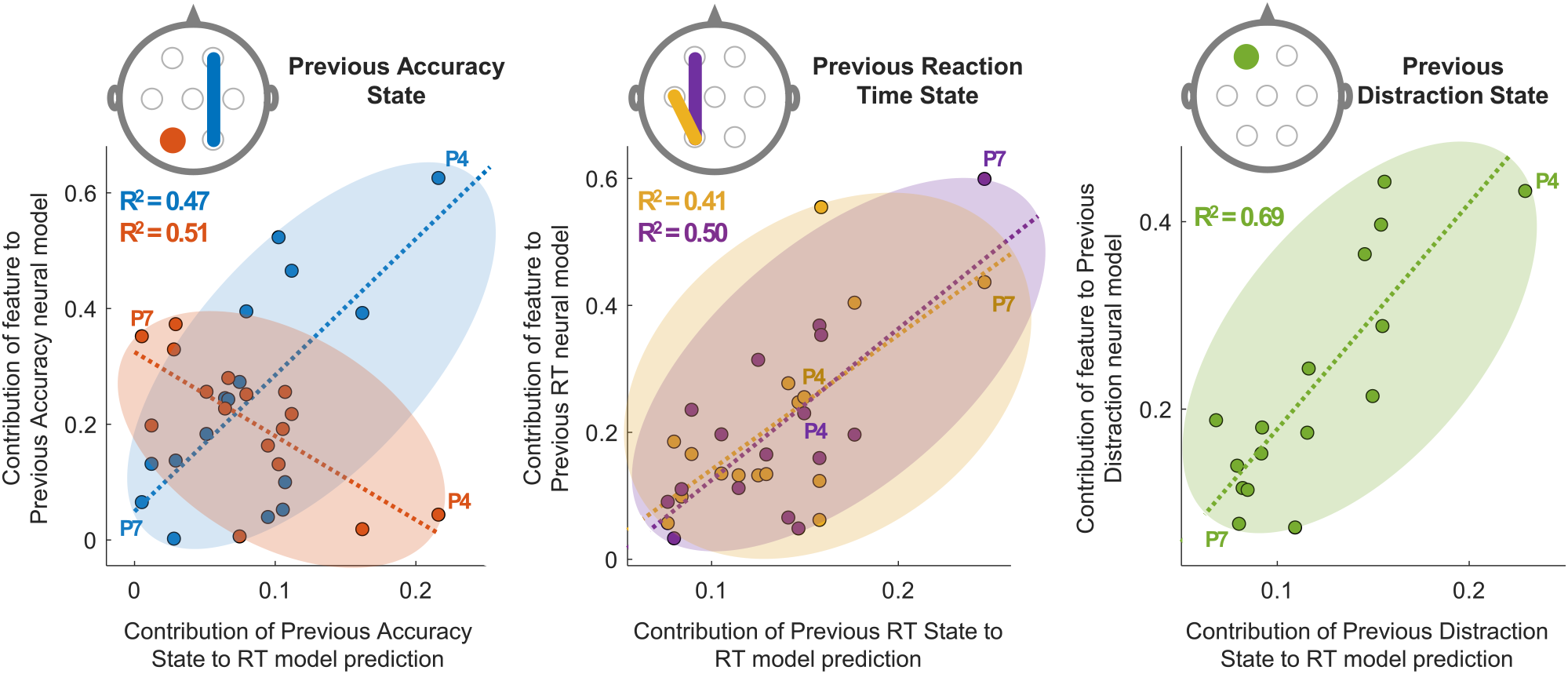
The highest linear relationships between the weights of the internal states for the behavioral model (*β_x_*) and the weights of the EEG features for the neural model (*γ*), for each of the three states. We show only the features that produced a *R*^2^ > 0.4. For each scatter plot, the color of the dots representing all 16 participants matches the color of the feature shown on the head illustration which are shown in the transverse plane and the hemispheres are not flipped. The dotted line shows the fitted regression line. All these EEG features are in the gamma band, except for the temporal – parietal connection (Previous RT State), which is in the alpha band.

Interestingly, for the Previous Accuracy State, we observed a different relationship for the left parieto – occipital gamma activity versus right fronto – parietal gamma connectivity. Overall, the more that participants actively remembered past errors, the more we observed right fronto – parietal gamma connectivity contributing to the prediction of the Previous Accuracy State, which weighed into RT. For example, P4 highly weighed previous accuracy in the behavioral model while P7 did not, and this was reflected in the weight of right fronto – parietal gamma connectivity in the neural model. The opposite trend occurred with the left parieto – occipital gamma activity. For P7, who barely weighed previous accuracy in predicting RT, the left parieto – occipital gamma weight from the neural model was high, while P4’s was low.

For the Previous RT State, the two strongest linear relationships were positive and found in the left fronto – parietal gamma and left temporo – parietal alpha connections. In other words, as the EEG functional connectivity of these two pairs of left-hemisphere regions weighed into the Previous RT State more, we observed participants were more likely to maintain their current trend of increasing or decreasing RT across trials.

Finally, the largest *R*^2^ value (0.69) observed reflected the relationship between the Previous Distraction State and left frontal gamma activity. For this state, since the correlation was positive, the more that participant’s RT was affected by distractors, the more that left frontal gamma activity weighed into predicting the Previous Distraction State.

## Discussion

The current study suggests a new perspective on behavioral and neuroimaging data related to experimentally controlled factors, and illustrates the importance of considering individual differences in dynamic internal states related to strategy, attention and motivation. Previous work on the present dataset (10) revealed that attentional capture by no-longer relevant information was associated with lower posterior alpha power, and that such capture was stronger when spatial relations rather than retinotopic locations were maintained in working memory, as indicated by both ERP magnitude and behavioral effects. The current results indicated, however, that this group effect of the experimental manipulation was small compared to variability in behavior and neural activity within and across individuals. This variability, however, was not noise. It was highly predictable. In particular, three states were found to be associated with task performance across individuals.

First, the Past Accuracy State reflected the influence of errors committed on previous trials. Our findings of right fronto-posterior couplings associated with this state are consistent with previous findings in which errors resulted in greater top-down, proactive control of information processing and response inhibition, mediated in part by right prefrontal cortex, in order to reduce the influence of irrelevant information (10, 13, 14, 19, 20). Following errors, trial-by-trial adjustments were implemented, more so by some participants than others, and appeared to be a major determinant of inter-individual variability in behavioral performance.

Second, the Previous RT State and its associated EEG activity appeared to reflect carry-over effects from one trial to the next regarding information processing, or perhaps attentional arousal and task engagement. Here, in contrast to the Past Accuracy state, left gamma frontoparietal and alpha parieto-temporal couplings were predominantly involved in predicting the Previous RT State and, thus, RT. Activity and connectivity in the alpha and gamma bands have previously been associated with working memory maintenance and information processing (21, 22). Since P4 and P7 displayed similar accuracy levels in the working memory task, but different degrees of association between this left hemisphere EEG connectivity and RT, the activity identified in the current analysis is likely not directly related to working memory processes. Further research will be needed, using source localization, to better understand the contribution of the Previous Reaction Time State and its associate neural activity.

Third, the Previous Distraction State referred to the distraction elicited by the incongruence between the hemifield of the distractor and the hemifield of the working memory test, relative to when the elements were congruent (i.e., displayed in the same hemifield). Our previous work (10) revealed a facilitating effect on task performance, on average in the group analysis, when the distractor and the subsequent working memory test were displayed in the same hemifield. Conversely, when the hemifields of the distractor and of the working memory test were different, decreased performance was observed. In the current study, the Previous Distraction State indicated the extent to which an individual’s performance on previous trials was affected by the congruence of the distractor and test stimuli. Additionally, the state was associated with left frontal gamma activity and predicted the extent to which a participant’s reaction time was affected by congruence on future trials. As with the other internal states, the degree to which the Previous Distraction State and its associated neural activity affected future RT was again highly variable across participants. Despite this variability, the Previous Distraction State coefficient was positive in all participants, consistent with the previous findings in the group analysis. The current analysis, however, adds additional information that is important to understand individual differences in this kind of distraction effect. The distractor color was relevant in a different task that the participants also performed. The Previous Distraction State indicates how much the participant’s distraction in previous trials predicts their RT in a current trial. That is, the contribution of the state indicates how much the individual’s tendency to be distracted by those stimuli consistently lingered from trial to trial. The left frontal activity, thus, could reflect the degree to which the participants were thinking about the rules of the tasks as it related to the distractors versus focusing only on the spatial working memory stimuli. Thus, again, the current analysis revealed factors affecting behavior and neural activity during the experimental task that are highly variable both within and across individuals. In this case the state may be related to strategy or meta-cognitive processes in order to stay focused on the current task.

Overall, the current study showed the promise of a dynamic systems modeling approach for understanding individual differences in cognitive control. A typical approach to understanding an individual’s cognitive strengths and weaknesses compares overall performance in a task that, for example, challenges selective attention versus overall performance in a task that challenges learning from feedback, or working memory capacity. The cognitive abilities of clinical populations have often been assessed in a similar manner. The method demonstrated here could provide a much more sensitive and specific measure of both trait-like and state-dependent cognitive abilities in both individuals and groups. For example, previous research has indicated that sensitivity to reward (or loss) appears to be a trait-like aspect of individual differences in cognitive performance, related to dopamine genotype (23). An individual’s sensitivity to reward and their control over reward-related behavior, however, can also vary within an individual across time based on moment-to-moment changes in various contextual factors (24). Susceptibility to distraction is also thought to be somewhat trait-like (25) although it can change with sleep deprivation (26), age (27), learning (28), and other factors. While many task performance measures have poor test-retest reliability (29), the coefficients identified using our modeling approach may be more reliable measures of individual differences, because they measure specific factors that feed into performance. Additionally, our dynamic systems modeling approach could also be applied in the same individuals across multiple tasks and across testing sessions. This would provide critical information to evaluate whether an individual who was, for example, more sensitive to distraction than their peers in a given task was consistently more sensitive across time and experimental contexts. The identification of neural features underlying susceptibility to distraction, or feedback sensitivity, etc., would also provide model-driven targets for training or neuromodulation interventions for improving performance (30, 31).

## Author Contributions

S.C. conceived and planned the experiments. T.H. carried out the experiments. C.B. and S.V.S. designed the model and the computational framework and analysed the data. All authors contributed to the interpretation of the results. C.B. took the lead in writing the manuscript. All authors provided critical feedback and helped shape the research, analysis, manuscript, and all approved the final manuscript version for submission.

## Acknowledgments

The authors would like to thank Kara Blacker and Brian Anderson for their work on the original experimental study and Michelle DiBartolo for her work developing the modeling approach using a different data set. This work was supported by the Johns Hopkins Science of Learning Research Grant and NIH/NINDS T32 NSO70201 (Interdisciplinary Training in Biobehavioral Pain Research).None of the authors have competing interests.

